# Expression biomarkers used for the selective breeding of complex polygenic traits

**DOI:** 10.1101/076174

**Authors:** M. Marta Guarna, Shelley E. Hoover, Elizabeth Huxter, Heather Higo, Kyung-Mee Moon, Dominik Domanski, Miriam E.F. Bixby, Andony P. Melathopoulos, Abdullah Ibrahim, Michael Peirson, Suresh Desai, Derek Micholson, Rick White, Christoph H. Borchers, Robert W. Currie, Stephen F. Pernal, Leonard J. Foster

## Abstract

We present a novel way to select for highly polygenic traits. For millennia, humans have used observable phenotypes to selectively breed stronger or more productive livestock and crops. Selection on genotype, using single-nucleotide polymorphisms (SNPs) and quantitative trait loci (QTLs), is also now applied broadly in livestock breeding programs; however, selection on protein or mRNA expression markers have not been proved useful yet. Here we demonstrate the utility of protein markers to select for disease-resistant behaviour in the European honey bees (*Apis mellifera* L.). Robust, mechanistically-linked protein expression markers, by integrating cis and trans effects from many genomic loci, may overcome limitations of genomic markers to allow for selection. After three generations of selection, the resulting stock performed as well or better than bees selected using phenotype–based assessment of this trait, when challenged with disease. This is the first demonstration of the efficacy of protein markers for selective breeding in any agricultural species, plant or animal.

**Significance statement:** The honey bee has been in the news a lot recently, largely because of world-wide die-offs due to the parasitic Varroa mite, which is becoming resistant to the chemical controls the bee industry uses. In this study, we show that robust expression biomarkers of a disease-resistance trait can be used, in an out-bred population, to select for that trait. After three generations of selection, the resulting stock performed as well or better than bees selected using the phenotypic best method for assessing this trait when challenged with disease. This is the first demonstration of an expression marker for selective breeding in any agricultural species, plant or animal. This also represents a completely novel way to select for highly polygenic traits.

## Introduction

European honey bees are a keystone species in agriculture as many crops depend on them for pollination and increased yield^4^. Honey bee colonies have been dying at increased rates over the past decade, largely due to increased pressure from diseases and pests^5^. Since these pests and pathogens are continually evolving resistance to the synthetic chemicals used to treat them, the most sustainable, long-term solution for bee health is the development of selective breeding programs that can enrich natural disease resistance mechanisms. However, selective breeding in *A. mellifera* is particularly challenging because most traits are expressed at the colony level^6^, and due to the haplo-diploid sex determination system in bees, challenges in storing germplasm^7^, the requirement for heterozygosity at the complementary sex determination locus^8^ that severely limits in-breeding, and the tendency for queens to mate with up to two dozen different drones. These factors mean that continual selection is required to maintain stock.

Bees do, however, have some effective disease-resistance traits, which also happen to be highly polygenic: one example is the social immunity function known as hygienic behaviour. Bees exhibiting hygienic behaviour are more efficient at removing dead, diseased, or dying brood from the hive^9, 10^, enabling them to resist or at least co-exist with pathogens such as American Fouldbrood (*Paenibacillus larvae*) or parasites such as *Varroa* mites (*V. destructor*) that would otherwise kill the colony. A closely related but distinct trait known as *Varroa*-sensitive hygiene enables bees to detect and disrupt the life cycle of reproductive female *Varroa* mites^11^. Both hygienic behaviour and *Varroa*-sensitive hygiene are heritable^12, 13^ and can therefore be selectively bred for; likewise, QTLs and SNPs have been linked to each behaviour^14, 15^, opening the door for their use in marker-assisted selective (MAS) breeding. Historically, genomic markers have been favoured for MAS because of their stability and reliability, whereas expression markers such as the levels of transcripts or proteins, are typically thought to be too variable and dependent on environment for use in MAS.

However, even a closely linked DNA feature may not be sufficient for MAS in honey bees; *A. mellifera* has one of the fastest recombination rates (~32 cM/Mb) known among animals, and this will rapidly break down inter-allele linkage through repeated rounds of meiosis^16^. On the other hand, causally-linked expression markers should be more robust to recombination since their presence is required for the trait, even though they have historically been thought to be too dependent on environment. Here we use a panel of protein markers identified through a multi-generational study^3^ to guide selective breeding of disease-resistance traits in honey bees through three generations. By the third generation, bee stocks selected through MAS were able to resist disease as effectively as bees raised through conventional selective breeding using standard field tests, with no detectable loss of other desirable traits such as honey production. This is the first successful use of expression markers for MAS that we are aware of.

## Results & Discussion

We have previously identified seven proteins in adult worker bees’ antennae whose expression is tightly linked to hygienic behaviour^3^(HB). To complete a comprehensive panel of markers for use in selective breeding, we added six more proteins derived from the same data, four that showed tight correlation with *Varroa*-sensitive hygiene and grooming behaviour, and two more also linked to hygienic behaviour. Of the latter, one was missing in an initial dataset and therefore failed to meet the inclusion criteria we used, and the other (Fig. 1a) had been ‘lost’ due to an unrealized change in accession number between protein database versions. These thirteen biomarkers were complemented by two ‘housekeeping’ proteins, α-spectrin and β-tubulin, that showed zero correlation with any behaviours (Supplemental Table 1) to serve as loading controls. The biomarker panel had originally been discovered through untargeted, data-dependent liquid chromatography-tandem mass spectrometry but a less stochastic detection method was required for scanning hundreds of samples. Therefore, multiple reaction monitoring (MRM) assays^17^ were developed for up to five peptides from each protein (Supplemental Table 1, Fig. 1b), with priority given to peptides we had observed in the discovery of these biomarkers^3^. The use of stable isotope-labelled standard peptides of known concentrations allowed quantitation of each peptide in protein extracts from worker bee antennae (Fig. 1c).

**Figure 1:**
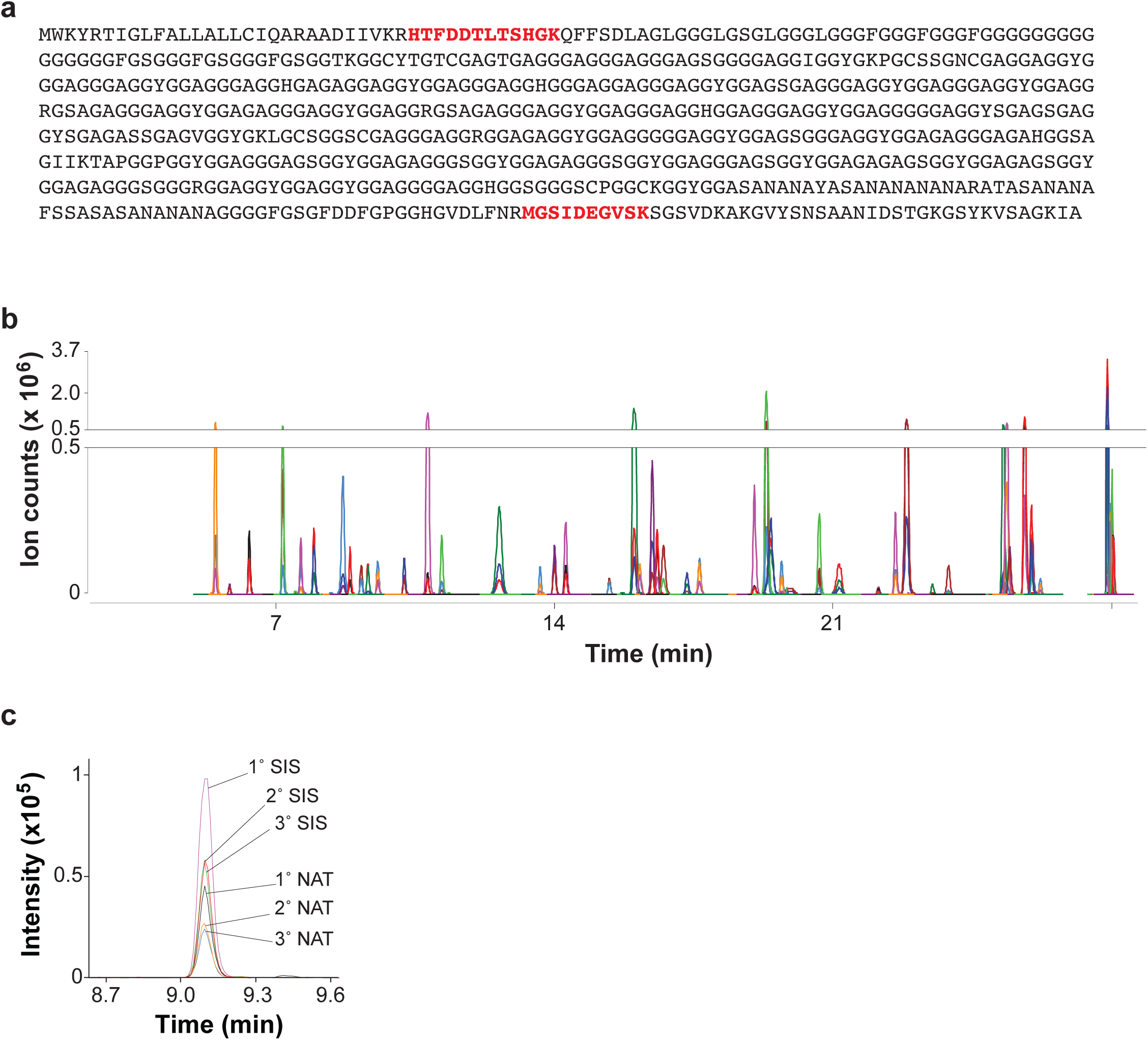
Multiple reaction monitoring assays for markers of disease resistance. a) amino acid sequence of gi:110761334, Glycine-rich cell wall structural protein-like protein, one of the markers of hygienic behaviour. The two peptides identified in the initial discovery are highlighted in red; these same two peptides were targeted with multiple reaction monitoring assays here. b) overlaid chromatograms of the three selected transitions for the stable isotope-labelled forms of all fifty-five peptides listed in Table 1 for the fifteen proteins comprising the biomarker panel. c) Transitions for the stable isotope standard (SIS) and natural (NAT) forms of MGSIDEGVSK from Glycine-rich cell wall structural protein-like protein. The primary (1˚) transition of each peptide was used for quantitation, while the secondary and tertiary transitions were used to confirm specificity.

To identify a robust initial breeding population of honey bees that was not already enriched in disease resistance behaviours, we surveyed the hygienic behaviour (HB) of 635 colonies from thirty-eight commercial beekeeping operations across western Canada in 2011. For this initial survey, we gave priority to beekeeping operations that bred their own bees so that the bees were well-adapted to the local climate^18^ and representative of stock being bred and used in Western Canada. All beekeepers donated or sold a subset of the tested queens to incorporate into the selection program. The hygienic behaviour scores in this initial survey varied regionally and ranged from 9.8% to 100%, with a median of 64% (Fig. 2a), matching levels of trait expression observed previously among unselected populations in a smaller survey within the same region^19^.

**Figure 2:**
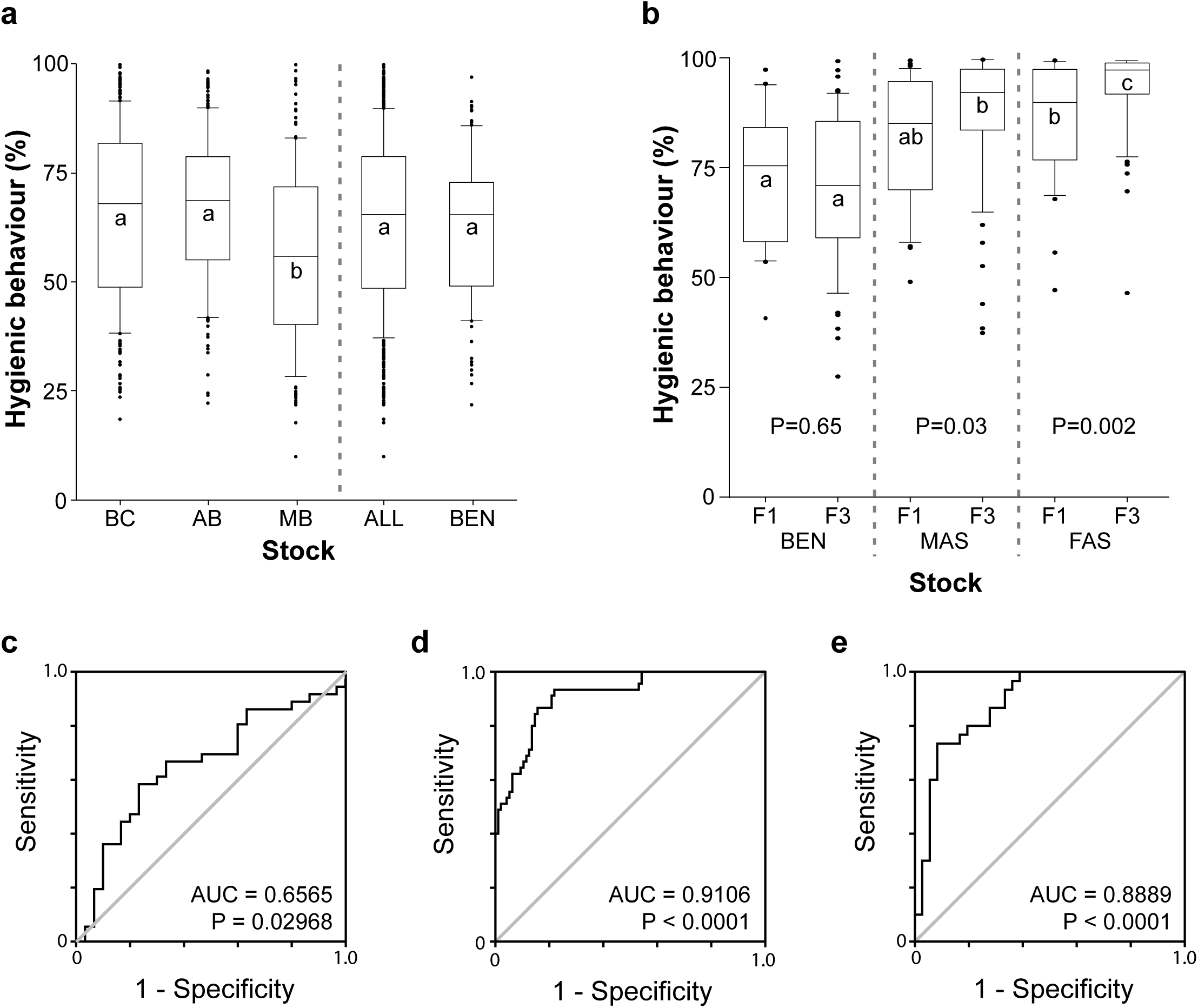
Starting distributions and enrichment of hygienic behaviour. BEN = benchmark, MAS = Marker-assisted selection, FAS = Field-assisted selection. (a) 90/10 box-and-whisker plots of the hygienic behaviour scores from all colonies in initial survey across Western Canada, in British Columbia (BC), Alberta (AB), Manitoba (MB)(left section) (Means followed by the same letter are not different from each other Tukey P<0.05), all colonies together (ALL), and the randomly selected starting benchmark population (BEN). ‘All’ is statistically identical to BEN (p=0.21, Analysis of Means Test)(b) The distribution of hygienic behaviour in the F1 and F3 generations of the benchmark population (left section, BEN, no statistical difference between F1 and F3, P=0.65, contrast), the colonies selected by the biomarker panel (middle section, MAS, F3 > F1, p=0.03, contrast), and the freeze-killed brood assay (right section, FAS, F3 > F1, p=0.002, contrast). Within each generation, means followed by the same letter are not different from each other Tukey P=0.05). Bottom: Receiver operating characteristics illustrating the performance of the F1 (c), F2 (d) and F3 (e) marker panels used for MAS.

For selection using markers, nurse bees from 468 of these colonies were dissected for quantitation of peptide markers in their antennae (Supplemental Table 2). From the colonies surveyed, two to four colonies were randomly selected from each beekeeping operation (for a total of 100 queens) to serve as benchmark, unselected stock (Fig. 2a): these (BEN) were statistically indistinguishable from the wider surveyed population (ALL). In addition, we selected queens from an additional 100 colonies, each with the highest HB scores in their apiaries. Their HB scores ranged from 38% to 100% with a mean score of 85%. We moved these queens to two breeding locations in Southern British Columbia and introduced them into colonies.

Over the next two years we reared three successive generations (F1 and F2 in 2012, F3 in 2013) from this initial population using the response of parental colonies in each of two ways: (1) the classic freeze-killed brood assay^9^ to quantify hygienic behaviour as a positive control (FAS, field-assisted selection), and (2) the levels of the best-performing subset of the peptide markers in Supplemental Table 3 (MAS, marker-assisted selection). For the selective breeding, the F1 and F2 queens were generated by instrumental insemination of virgin queens reared from the selected colonies using semen pooled from a random collection of drones from all the selected colonies in the appropriate stock. The same pooling of semen among selected was accomplished for F3 queens by closed breeding in isolated mountain valleys in southeastern British Columbia where no known drone sources existed. In addition to these selective breedings, benchmark stock (BEN) was maintained through unselected open mating, as a control.

The freeze-killed brood field assay (FAS) is the gold standard for identifying hygienic behaviour^9^ and colonies selectively bred using FAS showed the greatest enrichment of hygienic behaviour (Fig. 2b) over three generations. Notably, selective breeding based on the panel of protein markers (MAS) was also effective for enriching hygienic behaviour, demonstrating the potential of this technique in selective breeding.

By measuring hygienic behaviour in the colonies bred using MAS we could also monitor the specificity and sensitivity characteristics of the biomarker panels; while there was a statistically significant improvement detectable even in F1, by F2 there was a very marked improvement that was little changed in the F3 (Fig. 2c, d, e). The distribution of hygienic behaviour in the unselected BEN colonies, however, was unchanged between the starting group and the final population (Fig. 2b, left-most plot).

Hygienic behaviour can confer resistance to brood pests and pathogens that contribute to honey bee colony losses. We therefore evaluated how well colonies headed by selected queens performed under disease (American foulbrood (AFB), *P. larvae)* and mite (*V. destructor)* challenge conditions. Selected and benchmark stocks, as well as imported stock commonly used by Canadian beekeepers, were inoculated with either parasitic mites or AFB bacterial spores at levels that would normally result in high levels of colony mortality.

To test for the ability to survive with *Varroa* mites, in the early summer 2012 we pooled a large population of worker bees from colonies that were infested with *V. destructor* (about three hundred mites per colony, or a 3.5% infestation rate based upon mites per 100 bees) and aliquoted 8,600 bees (1 kg of bees) into individual colonies.

Twenty-three F3 selected or unselected queens were then randomly introduced into these individual colonies and the colonies were left untreated until the following spring; a fall survey for mites measured infestation rates of 23.0 ± 1.4 mites per 100 bees. For AFB, F3 colonies normalized for population received a frame with 225 cm^2^ of brood comb with 30 to 54% of wax cells showing visible *P. larvae* disease symptoms on each side. We assessed the impact of the parasite and pathogen challenge on winter survival for the varroa challenge and for overall survival of asymptomatic colonies for the AFB challenge (Fig. 3).

**Figure 3:**
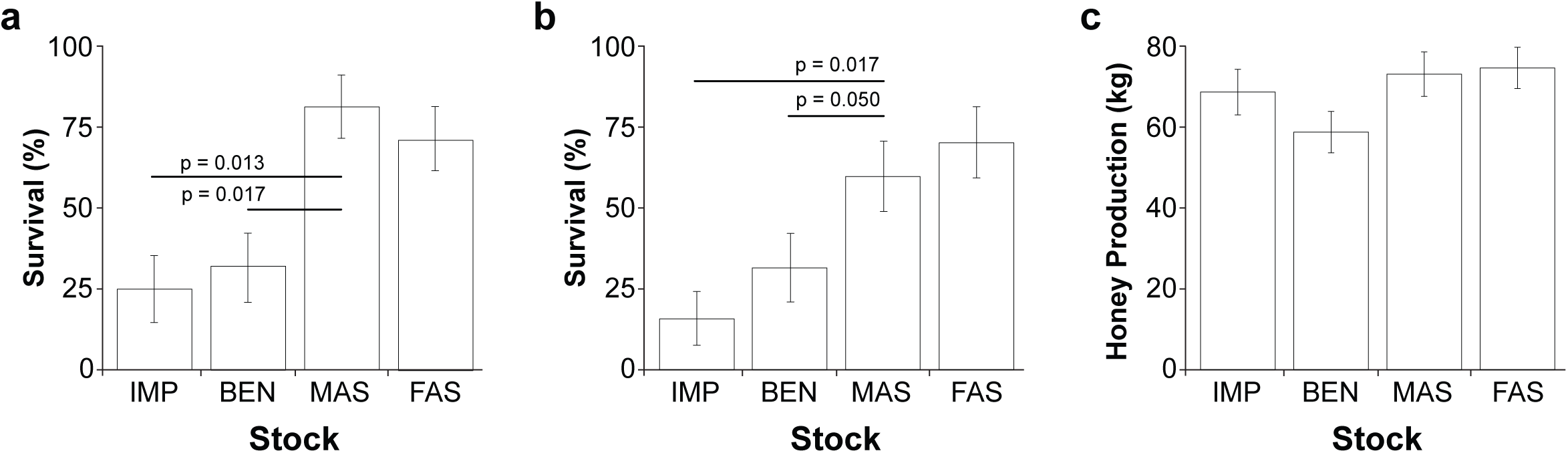
Performance of selected stock. IMP = imported stock, BEN = benchmark, MAS = Marker-assisted selection, FAS = Field-assisted selection. (a) Difference in winter survival of F3 generation colonies headed by queens from each stock type that were challenged with *Varroa* mites (*Varroa* challenge) (d.f. 3, Chi Sq 14.84 p > chi = 0.002). (b) Difference in symptom-free survival when challenged with American Foulbrood (*P. larvae*; AFB challenge:) (d.f. 3, Chi Sq 12.65 p > chi = 0.0054). Horizontal lines represent Holm-Bonferonni adjusted single degree of freedom contrasts between MAS selected stock and the benchmark and imported stock controls. Siimilar results were found for FAS, with FAS survival higher than the BEN and IMP stocks for both the *Varroa* challenge (p=0.05 and p=0.025, respectively) and AFB challenge experiments (p=0.025 and p=0.013, respectively). Error bars represent the standard error of the binomial proportion. (c) Honey produced per colony for all stocks tested at three experimental sites in Alberta and Manitoba. There was no significant difference in honey production among the four stocks tested (d.f. 3,161; F=2.12, p=0.099).

To check that an independent but critical performance indicator of these stocks was not being degraded by too much focused selection on another trait, we also examined the ability of F3 colonies to collect honey in a separate study from the disease-challenge experiments. Reassuringly, honey production was not affected by selection for hygienic behaviour, regardless of which selection method was used (Fig. 3c).

The results of these experiments were subsequently used to model the economic impact of integrating the use of marker-selected stock into a Canadian beekeeping operation. The marker-selected colonies’ increased disease resistance and greater survival rates enhanced beekeeper profit. The economic modeling shows that when a beekeeper replaces his conventional colonies with MAS colonies selected for hygienic behavior we see up to a 5% increase in profit for a 40-colony apiary. This is likely an underestimate as we have not assumed any increase in colony productivity for the disease-resistant MAS colonies. The greatest economic value derived from MAS colony adoption was when resistance to traditional Varroa treatments was modeled in the apiary. To reduce the risk of economic loss associated with treatment resistance (or an equivalent ineffective/ lack of treatment), MAS colonies replaced 25%, 50% and 100% of traditional colonies within the apiary showing profit increases of over 300%, 600% and 800% respectively compared to the no-MAS apiary.

The first genetic modification of bees has been reported^21^ but industrial use of genetically modified bees is unlikely to be accepted by the public at this time. Therefore, tools that enable accelerated stock enrichment without resorting to genetic modification are highly desirable. While genetic markers for hygienic behaviour and *Varroa*-sensitive hygiene have been identified^14, 15^, they are unlikely to be linked tightly enough to be robust to the high recombination rate bees exhibit for more than a few generations. In addition, there may be several loci missing since both traits are highly polygenic^22^. Expression markers integrate many different *cis-* (e.g., transcriptional enhancer elements) and *trans-* (e.g., transcription factors) effects so if they are functionally linked to the trait in question then they should be robust to recombination. We have not yet shown that the markers we use here are functionally linked to hygienic behaviour, *Varroa*-sensitive hygiene, or grooming but they are tightly linked enough to enrich the trait as quickly as the best conventional methods available.

The most important pests and pathogens of bees are currently controlled with acaricides, antibiotics and antimycotics, but emerging resistance to these treatments may be partially responsible for the higher level of colony losses seen over the past seven years^23^. These exogenous treatments can leave residues in the hive^24^ and honey though so disease-resistant stock is seen as the best solution, simultaneously reducing colony losses and the need for synthetic chemicals while ensuring food safety. Marker-assisted selection of honey bees will enable more sophisticated breeding of this critical agricultural service provider.

Selective breeding is a vital tool for improving yields and disease resistance in all plants and animals used in agriculture. Marker-assisted selection has the potential to be more precise and more robust to external influences; it has been widely used in certain plants^25^ and animals^26^. To date, however, the markers used have been genomic loci exclusively, starting with restriction fragment length polymorphisms and leading up to single nucleotide polymorphisms. This is undoubtedly due, in part, to the availability of efficient genetic approaches for finding such markers. It is also a matter of focus though: researchers have spent more time looking for genetic loci than for expression markers (i.e., transcripts or proteins) because the latter have been hitherto considered to be too dependent on environment. Here we have shown that expression markers can be used to select for a very complex, polygenic trait. Even in this proof-of-principle with a first-generation panel of markers, MAS was as efficient at enriching disease-resistance as FAS methods: bees bred using marker-assisted selection could resist levels of disease that would typically kill 75% or more of unselected colonies. The data presented here have implications beyond bees: this is the first demonstration of marker-assisted selection in livestock using expression markers and it opens the door for molecular diagnostic approaches for selecting complex polygenic traits that are recalcitrant to genetic mapping methods^27^.

## Acknowledgements

The authors wish to thank Susan Cobey for assistance with instrumental insemination, Dominik Domanski and Derek Smith for MRM assay development, and Immacolata Iovinella and Paolo Pelosi for providing recombinant Odorant Protein 16 for MRM peptide selection. We also thank Lisa Babey, Zoe Rempel, Rasoul Bahreini, Lindsay Geisel, Carl To, Jaclyn Deonarine, Daryl Wright, Cole Robson-Hyska, Sarah Carson, Dave Holder, and Lynae Ovinge for technical assistance. We sincerely thank each cooperating beekeeper that facilitated hygienic behaviour testing in their operations and donated queens to the project. This work was supported by funding from through the BeeIPM project (107BEE).

## Author Contributions

LJF, SFP, RWC, and MMG conceived the experiments. LJF, MMG, SFP, RWC, SEH, EH, AI, APM, and HH designed the experiments. EH and HH managed the selective breeding. RW and MMG developed the statistical treatment of the biomarkers and refined the prediction models. KMM oversaw the proteomic sample collection and processing. SEH, SFP, MMG and KMM analysed data from freeze-killed brood and MRM assays data to select breeding colonies. All authors except RW, MB, DD and CB helped with sample collection, hygienic behaviour testing, and general beekeeping activities. DD and CB developed and applied the multiple reaction monitoring assays. SEH, AI, MP, SD, DM, SFP and RWC conducted the *Varroa-* and *P. larvae*-challenge experiments, as well as the evaluation of honey production. MEFB developed the economic model.

